# CDKL5 deficiency impairs TBK1-mediated autophagy and clearance of neuronal protein aggregates

**DOI:** 10.64898/2026.09.13.751248

**Authors:** Zhongju Zou, Bilal Kahn, Salwa Sebti, Jason Lucavs, Giomar Rivera-Cancel, Josephine Thinwa

## Abstract

The clearance of unwanted protein aggregates is essential for maintaining proteostasis and cellular function, particularly in long-lived cells such as neurons, yet the signaling pathways that activate selective autophagy of protein aggregates remain incompletely understood. Here, we identify the neurodevelopmental kinase CDKL5 as an upstream regulator of a signaling pathway involving the TBK1 adaptor SINTBAD and the selective autophagy receptors p62 and TAX1BP1. CDKL5-deficient mice show age-dependent accumulation of detergent-insoluble protein aggregates in the brain, accompanied by impaired TAX1BP1 recruitment and reduced p62 Ser405 phosphorylation. In cultured cells and primary cortical neurons, loss of CDKL5 delays clearance of puromycin-and proteasome-inhibitor-induced aggregates in a manner dependent on CDKL5 kinase activity. Mechanistically, CDKL5 kinase activity is required for SINTBAD Ser504 phosphorylation, a SINTBAD modification that promotes TBK1 activation, resulting in p62 Ser403/405 phosphorylation and TAX1BP1-dependent aggregate clearance. Phosphomimetic SINTBAD rescues these responses in CDKL5-deficient cells. These findings define a CDKL5/SINTBAD/TBK1 signaling axis that couples proteotoxic stress to activation of selective autophagy receptors and identify impaired proteostasis as a previously unrecognized consequence of CDKL5 deficiency.

## Introduction

Mechanisms of protein quality control maintain cellular homeostasis through the removal of aggregated proteins. Key to proteostasis is autophagy, an evolutionarily conserved mechanism in which large macromolecules such as damaged organelles and protein aggregates become sequestered into double-lipid membrane vesicles destined for lysosomal degradation^1^. Impaired autophagy has particularly been linked to the pathogenesis of neurodegenerative disorders such as Alzheimer’s and Parkinson’s diseases, which are associated with the aberrant accumulation of protein aggregates such as amyloid and tau^2^. The long-lived, highly metabolic, and non-renewable nature of neurons makes them highly dependent on autophagy for the maintenance of homeostasis^3–5^. To avoid proteotoxic stress, neurons maintain tight quality control of proteins by balancing biogenesis, folding, and degradation^3,6^.

The selective clearance of protein aggregates, termed aggrephagy, depends on a coordinated set of autophagy receptors that recognize ubiquitinated cargo and couple it to the core autophagy machinery^7^. Receptors such as p62/SQSTM1 and NBR1 bind ubiquitinated proteins and promote their condensation into larger inclusions, concentrating aggregated proteins for subsequent degradation^8^. Efficient clearance, however, requires more than cargo recognition: TAX1BP1 subsequently couples these condensates to the autophagy initiation machinery, in part by recruiting the kinase TBK1 through adaptor proteins including SINTBAD and NAP1^9^. Activated TBK1 phosphorylates selective autophagy receptors, including p62 at Ser403/405, enhancing binding to ubiquitinated cargo, autophagic sequestration and degradation of ubiquitinated cargo^10,11^. Thus, aggrephagy is not simply a consequence of aggregate recognition but requires regulated signaling events that promote progression from cargo condensation to autophagosome engulfment of cargo^9^. The upstream pathways that sense proteotoxic stress and activate this receptor network remain incompletely understood.

The importance of autophagy extends beyond the maintenance of proteostasis in aging neurons. Pathogenic human variants affecting core autophagy proteins, including ATG5 and ATG7, cause severe childhood neurodevelopmental disorders, demonstrating that appropriate autophagic activity is also required for normal nervous system development and function^12,13^. In our previous work, we identified a neurodevelopmental kinase called cyclin-dependent kinase-like 5 (CDKL5), as a regulator of virophagy, the selective autophagic clearance of viral components during infection^14^. Loss-of-function variants in CDKL5 cause CDKL5 deficiency disorder (CDD), a devastating neurodevelopmental epileptic encephalopathy characterized by early-onset seizures and profound neurological impairment^15^. Despite its importance in human disease, the molecular pathways controlled by CDKL5 remain incompletely defined^16^. Our identification of CDKL5 as a regulator of selective autophagy raised the possibility that its function extends beyond antiviral defense to more general mechanisms of protein quality control.

Here, we identify CDKL5 as a regulator of aggrephagy and define a signaling pathway through which it promotes the clearance of aggregation-prone proteins. Using cell culture, primary neurons, and aged mouse brains, we show that loss of CDKL5 impairs the clearance of multiple forms of proteotoxic cargo and leads to the progressive accumulation of detergent-insoluble ubiquitinated proteins in vivo. Mechanistically, CDKL5 acts upstream of the SINTBAD–TBK1 signaling axis, phosphorylating SINTBAD to promote TBK1 activation and coordinate the activities of the selective autophagy receptors TAX1BP1 and p62. These findings identify CDKL5 as an upstream regulator of the signaling machinery that couples proteotoxic stress to selective autophagy and suggest that defective protein quality control may contribute to the neurological consequences of CDKL5 deficiency.

## Results

### CDKL5 is crucial for mitigating age-related protein aggregate accumulation in the brain

We have previously shown CDKL5 is important for virus-induced autophagy^14^; however, whether CDKL5 plays a larger role in autophagy and maintenance of cellular proteostasis is unknown. We sought to further investigate if in vivo, CDKL5-deficient mice exhibit increased protein aggregates in the brain. Mechanisms that maintain cellular proteostasis in the brain (including autophagy) wane with age leading to an accumulation of protein aggregates made of molecules such as lipofuscin^4^. Thus, we compared young adult (4-month-old) wild-type (WT) and CDKL5 KO mice with groups aged 18 or 24 months to assess whether loss of CDKL5 impacts age-dependent accumulation of protein aggregates. Brain sections were stained with ProteoStat, a fluorescent dye that labels protein aggregates. No differences were detected in the hippocampal or cortical regions of young mice; however, CDKL5 KO mice exhibited significantly increased aggregates in the cortex at 18 months and in both the hippocampus and cortex at 24 months compared with age-matched wild-type controls (Fig.1 A-C). Western blot analysis also showed increased high-molecular weight ubiquitinated proteins in the hippocampus and cortex of CDKL5 deficient mice (Fig. 1D).

**Figure 1:**
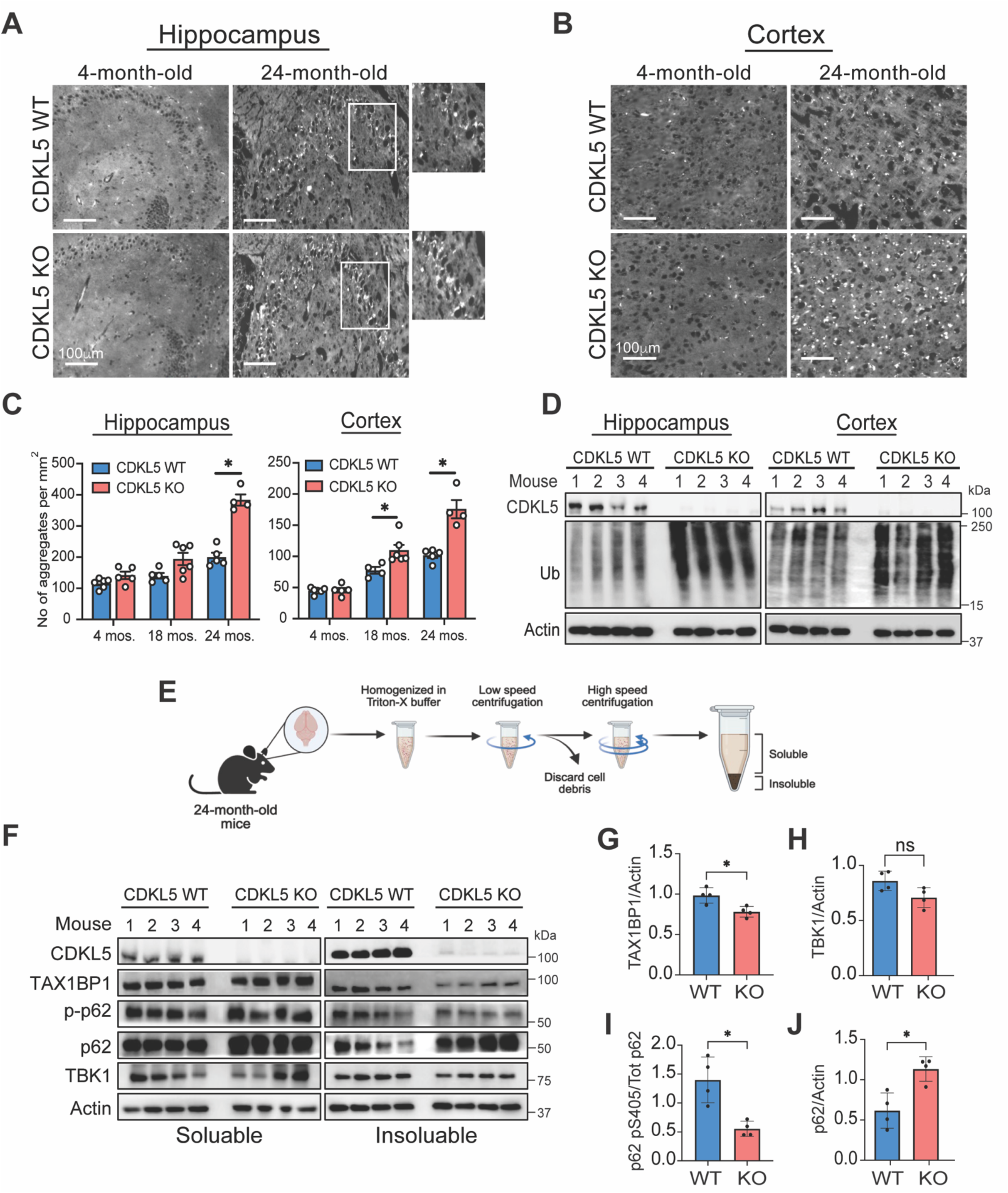
CDKL5-deficient mouse brains accumulate protein aggregates with aging. Young (4-month-old) and old (24-month-old) WT and CDKL5 KO mouse brains were harvested (n=4-5 males per group). A-B) Staining for protein aggregates with ProteoStat in the A) hippocampus and B) cortex, with C) quantification of the number of punctate aggregates per mm². Scale bar represents 100 µm. D) Western blot analysis of brain lysates for detection of ubiquitinated proteins. E) Schematic of the sequential detergent extraction protocol used to separate Triton-soluble and - insoluble protein fractions from brain tissue. F) Western blot analysis of Triton-soluble and - insoluble brain lysates (n=4 mice per group). G-J) Densitometry quantification of the designated proteins. All histobars represent the mean ± SE. Statistical analysis was performed using two-way ANOVA with Sidak’s correction for multiple comparisons (C) and Mann-Whitney U test (G-J).

To identify specific proteins within the aggregates, we performed high-speed centrifugation of brain lysates from 24-month-old mice to separate triton-soluble and insoluble fractions, as aggregated proteins sediment into the insoluble fraction (Fig. 1E). CDKL5 was detected in both the soluble and insoluble fractions, suggesting its incorporation into protein aggregates. We next assessed levels of two key autophagy receptors, TAX1BP1 and p62, which are essential for the selective degradation of ubiquitinated proteins^8,17^. p62 mediates the initial sequestration of ubiquitinated proteins into large condensates, whereas TAX1BP1 incorporates into the condensates to recruit the autophagy initiation machinery^8,9^. In CDKL5-deficient brain lysates, p62 showed significant accumulation in the insoluble pellet, while TAX1BP1 recruitment was significantly reduced compared with wild-type controls (Fig. 1F, 1G, and 1J). We next examined whether p62 within the aggregates was phosphorylated at Ser405 (human Ser403), a modification known to depend on TAX1BP1 and the kinase TBK1 for the clearance of protein aggregates^9^. Although TBK1 levels were not significantly different between WT and KO mice, the amount of phosphorylated p62 relative to total p62 was significantly reduced in the absence of CDKL5 (Fig. 1F, 1H, and 1I) overall suggesting a potential defect in autophagy receptor recruitment and function in CDKL5-deficient brains.

### CDKL5-deficient cells have impaired clearance of protein aggregates

Given the accumulation of protein aggregates observed in vivo in CDKL5 KO brains, we next used in vitro systems to determine whether CDKL5 plays a role in aggrephagy. To do this, we assessed autophagic clearance of puromycin-induced aggresome-like induced structures (ALIS), which are protein aggregates known to be cleared via autophagy^17,18^, in CDKL5-deficient HeLa cells. After 2 hours of puromycin treatment, WT, CDKL5 KO, and CDKL5 siRNA-knockdown (KD) cells all showed robust formation of ubiquitin-positive foci (Fig. 2A-B; Supp Fig. 1A-B). We then washed off the puromycin and monitored clearance of these ubiquitin foci over 6 hours as a readout of autophagic flux. CDKL5 KO and KD cells cleared ubiquitin foci at a significantly slower rate than WT cells (Fig. 2A-B; Supp Fig. 1A-B). Consistent with impaired autophagic degradation, blocking autophagic flux with Bafilomycin A1 (Baf A1) restored ubiquitin foci in non-coding siRNA-treated cells but had no additional effect in CDKL5 KD cells (Supp Fig. 1A-B). Immunoblotting for puromycin-labeled proteins similarly showed reduced clearance over time in CDKL5 KO cells (Fig. 2C).

**Figure 2:**
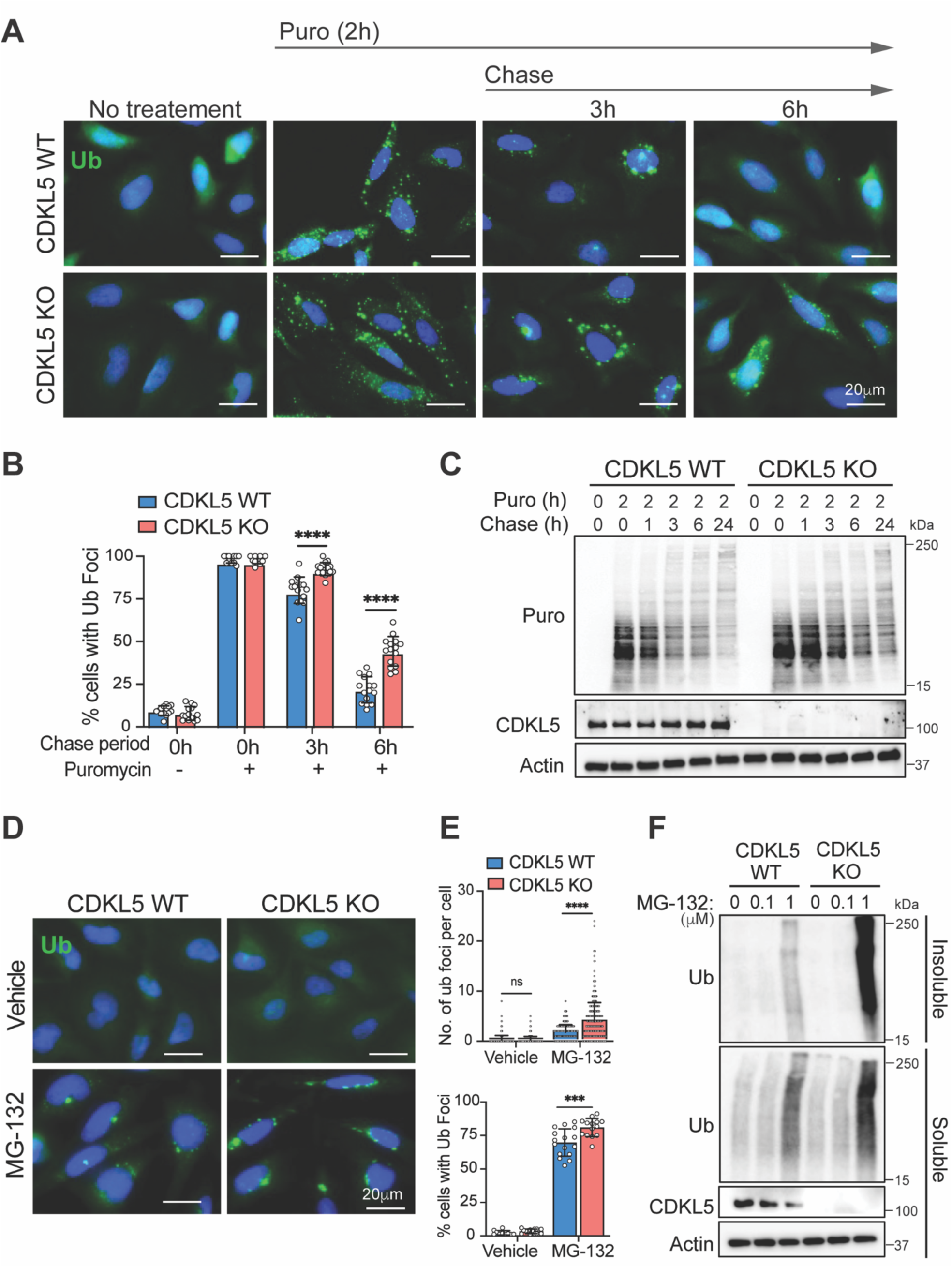
Loss of CDKL5 delays clearance of aggregated proteins. A) Representative fluorescence micrographs of ubiquitin in WT and CDKL5 KO HeLa cells treated with 5 μg/mL puromycin for 2 hours, washed, and replaced with fresh media for the designated chase period, with B) quantification of the percentage of cells with ubiquitin foci from 15 images and over 150 cells per condition across 3 independent experiments. Histobars represent the mean ± SE. Statistical analysis was performed using one-way ANOVA with Sidak’s correction for multiple comparisons. C) Representative western blot of puromycin-treated WT and CDKL5 KO cells over a 24-hour chase period, detecting puromycin, CDKL5, and actin. D) Representative fluorescence micrographs of ubiquitin in WT and CDKL5 KO HeLa cells treated with 1 μM MG-132 for 18 hours, with E) quantification of ubiquitin foci per cell and percentage of cells with ubiquitin foci. Histobars represent the mean ± SE of 15 images and over 150 cells per condition across 3 independent experiments. Statistical analysis was performed using one-way ANOVA with Sidak’s correction for multiple comparisons. F) Western blot analysis of WT and CDKL5 KO cells treated with increasing concentrations of MG-132 for 18 hours.

To further validate these findings, we treated cells with the proteasomal inhibitor MG-132 to induce misfolded protein accumulation and aggregation. CDKL5-deficient cells showed approximately twofold more ubiquitin foci and a higher percentage of foci-positive cells compared to controls (Fig. 2D-E; Supp Fig. 1D-E). Fractionation of cell lysates into detergent-soluble and -insoluble fractions revealed that CDKL5-deficient cells accumulated substantially more ubiquitinated protein in the insoluble fraction (Fig. 2F; Supp Fig. 1C), corroborating the puromycin data.

### CDKL5 facilitates clearance of aggregation-prone huntingtin protein

Autophagy is known to mediate the clearance of polyglutamine (polyQ) repeat expansion aggregate-prone proteins that cause cytotoxicity in disorders such as Huntington’s disease^19^. To determine if CDKL5 impacts the clearance of polyQ protein, we overexpressed a protein containing pathogenic expanded glutamine tracts expressed in exon 1 of the huntingtin-encoded gene (HttQ74-EGFP) or a shorter non-pathogenic form (HttQ23-EGFP) in WT or CDKL5 KO HeLa cells. Using a filter assay, we analyzed levels of large protein aggregates on western blot and detected significantly increased polyQ74 aggregates in CDKL5 KO cells compared to WT (Supp Fig. 2A-B). To validate these findings, HeLa cells expressing HttQ74-EGFP were treated with increasing concentrations of CDKL5 inhibitor (CAF-382)^20^. The inhibition of CDKL5 activity was assessed by detecting EB2 phosphorylation at Ser222, a modification that is well known to be mediated by CDKL5^21^. Levels of polyQ74 aggregates increased in a dose-dependent manner (Supp Fig. 2C).

### CDKL5 kinase activity is required for protein aggregate clearance

We next asked whether CDKL5 kinase activity is required for its role in aggrephagy by reconstituting CDKL5 KO cells with a kinase-dead mutant, CDKL5^K42R^, which is unable to effectively bind ATP^20,22^. Following puromycin treatment, CDKL5 KO cells expressing CDKL5^K42R^ cleared ubiquitin puncta at a rate similar to KO cells expressing empty vector (EV). In contrast, expression of WT CDKL5 significantly increased the rate of ubiquitin foci clearance, an effect that was reversed by Bafilomycin A1 treatment (Fig. 3A-B). To further test the requirement for CDKL5 kinase activity, we treated CDKL5 KO cells expressing EV, WT CDKL5, or CDKL5^K42R^ with MG-132. After 18 hours of treatment, cells expressing EV or CDKL5^K42R^ showed comparable and significant accumulation of ubiquitin foci by immunofluorescence, as well as increased insoluble ubiquitinated proteins by immunoblotting, relative to cells expressing WT CDKL5 (Fig. 3C-E). Together, these data indicate that CDKL5 kinase activity is required for its function in aggrephagy.

**Figure 3:**
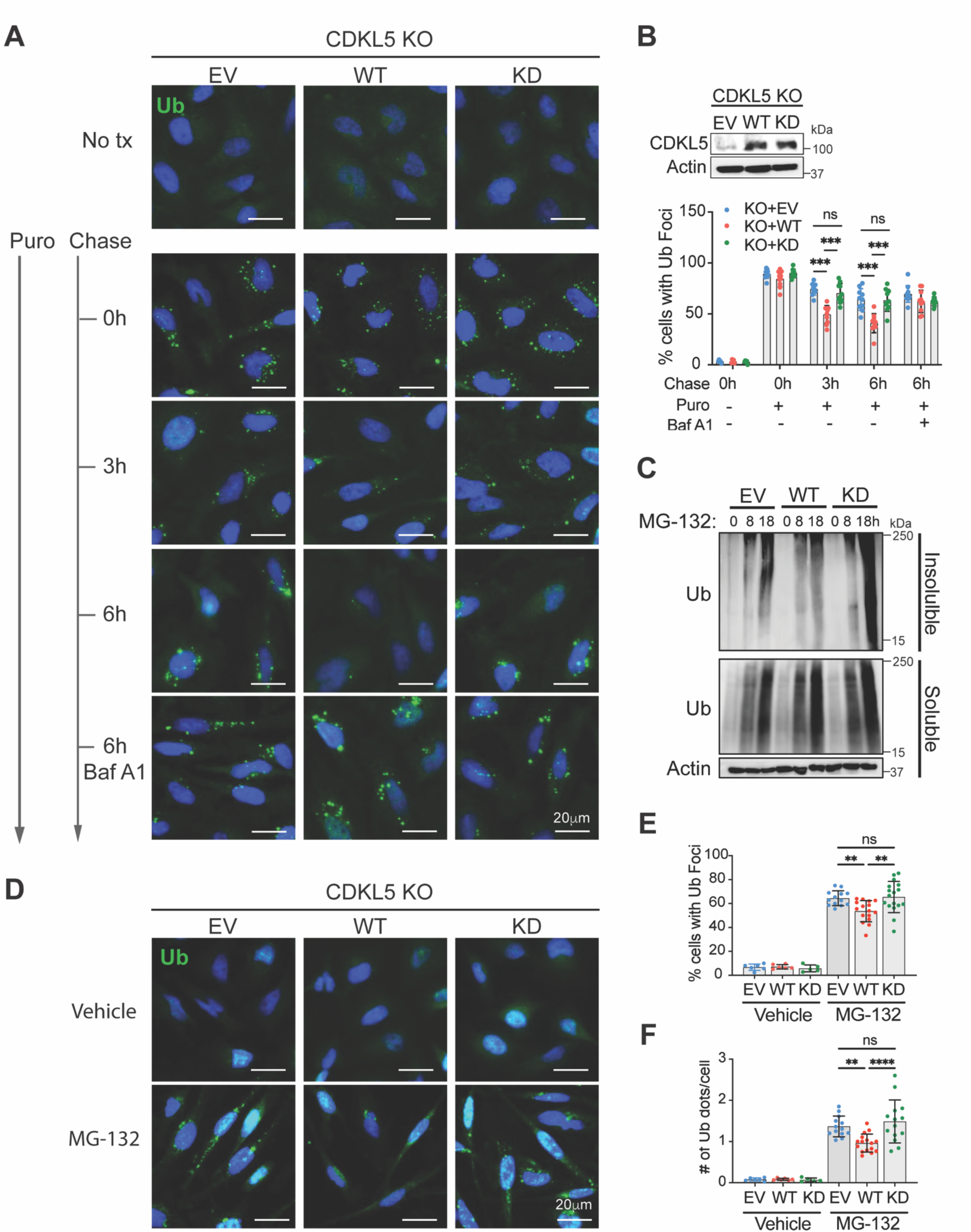
CDKL5 kinase activity is required for protein aggregate clearance. A) Representative fluorescence micrographs of ubiquitin in CDKL5 KO HeLa cells reconstituted with empty vector (EV), kinase-dead K42R mutant (KD), or WT CDKL5, treated with 5 μg/mL puromycin for 2 hours, followed by a post-wash chase period in the presence or absence of Bafilomycin A1 (200 nM). B) Western blot of CDKL5 expression in reconstituted KO cells, and quantification of the percentage of cells with ubiquitin foci. Bars represent the mean ± SE of triplicate samples with at least 100 cells; similar results were observed in three independent experiments. C) Western blot analysis of CDKL5 KO cells expressing EV, KD, or WT CDKL5, treated with MG-132 (1 μM) over a time course. Lysates were fractionated into Triton-soluble and -insoluble fractions. D) Representative fluorescence micrographs of ubiquitin in CDKL5 KO HeLa cells reconstituted with EV, KD, or WT CDKL5, treated with 1 μM MG-132 for 18 hours, with quantification of E) the percentage of cells with ubiquitin foci and F) the number of ubiquitin foci per cell. Bars represent the mean ± SE of triplicate samples with at least 100 cells across three independent experiments. Statistical analysis for B, E, and F was performed using one-way ANOVA with Sidak’s correction for multiple comparisons. **p < 0.005, ***p < 0.0005, ****p < 0.00005.

### TAX1BP1 and p62 autophagy receptors are impaired in CDKL5 deficiency

Our in vivo data suggested a potential dysfunction in TAX1BP1 and p62, which are known to be essential for capturing ubiquitinated protein aggregates and recruitment of autophagic machinery^8,17,23^. We therefore investigated if CDKL5 deficiency impacted their association with ubiquitinated cargo in vitro through immunofluorescence. Two hours of puromycin treatment increased TAX1BP1-positive foci in WT cells but not in CDKL5 KO cells (Fig 4A-B). After a 6-hour chase period post puromycin wash-out, these TAX1BP1/ubiquitin foci cleared significantly faster in WT cells compared to KO cells (Fig. 4 A-B). Notably, addition of BafA1 in WT HeLa cells restored TAX1BP1/ubiquitin foci but not in CDKL5 KO cells (Fig. 4A-B), suggesting that clearance of these foci in WT cells was autophagy-dependent. Conversely, p62/ubiquitin foci increased similarly in both WT and CDKL5 KO cells following puromycin treatment (Fig. 4C-D), but cleared significantly faster in WT cells after puromycin washout, and were restored to a greater degree by lysosomal blockade in WT cells relative to KO cells (Fig. 4C-D). Taken together these data suggest that CDKL5 has a greater impact on the recruitment of TAX1BP1 to ubiquitin foci compared to p62. We subsequently performed time-lapse live-cell imaging to track the movement of TAX1BP1 foci following puromycin treatment, and found that TAX1BP1 foci collided, fused, and cleared more rapidly in WT cells than in CDKL5 KO cells (Supplemental Fig. 3A-D).

**Figure 4:**
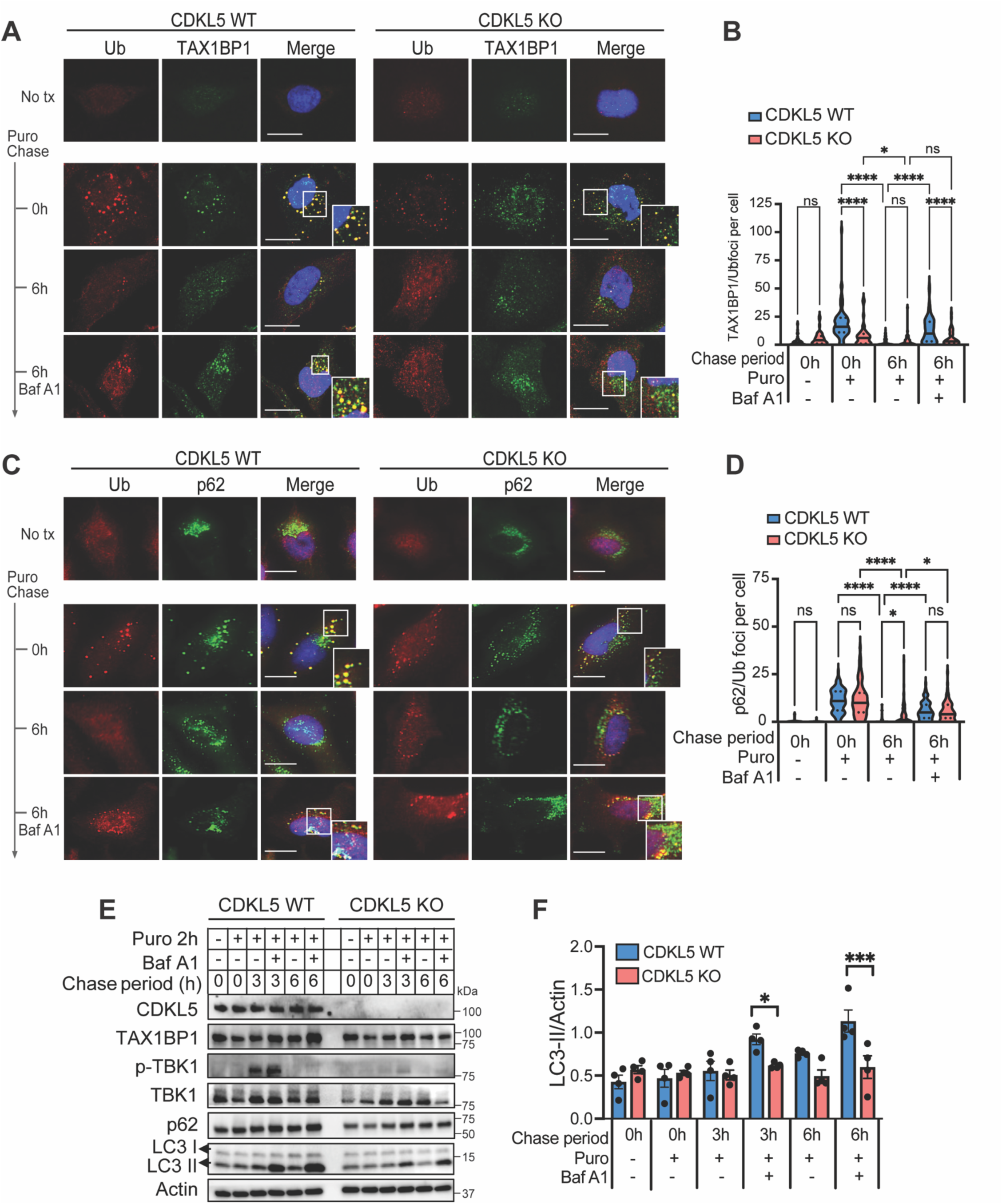
Impaired autophagy adaptor function in CDKL5-deficient cells. CDKL5 WT and KO HeLa cells were treated with puromycin for 2 hours, followed by a post-wash chase period of 6 hours in the presence or absence of Bafilomycin A1 (200 nM). Representative fluorescence micrographs of A) ubiquitin and TAX1BP1, with B) quantification of TAX1BP1-positive ubiquitin foci per cell, and C) ubiquitin and p62, with D) quantification of p62-positive ubiquitin foci per cell. Histograms represent over 50 cells per condition across 3 independent experiments. E) Representative western blot of puromycin-treated WT and CDKL5 KO cells, with F) LC3-II normalized to actin densitometry analysis from 3 independent experiments. Statistical analysis for B) and D) was performed using one-way ANOVA with Sidak’s correction for multiple comparisons, and for F), two-way ANOVA with Sidak’s correction for multiple comparisons. *p<0.05, **p < 0.005, ***p < 0.0005, ****p < 0.00005.

These findings suggested that CDKL5 may regulate an upstream signaling pathway that coordinates selective autophagy receptor function. TBK1 is a key regulator of selective autophagy that functionally interacts with both TAX1BP1 and p62 and undergoes trans-autophosphorylation at Ser172 to become catalytically active^9,24,25^. We performed immunoblotting to assess TBK1 phosphorylation and found that Ser172 could be detected as early as 3 hours after puromycin washout in WT cells but failed to be induced in CDKL5 KO cells (Fig. 4E). Notably, in the presence of BafA1, CDKL5 KO cells showed significantly less LC3-II accumulation compared to WT cells at both the 3-and 6-hour chase time points following puromycin washout, suggesting that loss of CDKL5 attenuates autophagic flux (Fig. 4E-F).

### CDKL5 deficiency in neurons impairs aggrephagy adaptor function

CDKL5 is highly expressed in neurons^26,27^. To investigate whether CDKL5 deficiency disrupts neuronal aggrephagy, we quantified the percentage of WT and CDKL5 KO mouse primary cortical neurons containing ubiquitin-positive protein aggregates following treatment with MG-132. CDKL5 KO neurons accumulated significantly more ubiquitinated proteins by 8 hours and cleared these proteins more slowly by 18 hours compared to WT neurons (Fig. 5A-B). Because autophagy adaptor proteins are themselves degraded as autophagosome substrates, their turnover can be monitored alongside that of ubiquitinated protein during MG-132 treatment, providing an additional readout of autophagic flux^28^. In CDKL5-deficient neurons, levels of p62, TAX1BP1, and high molecular weight ubiquitinated proteins all declined more slowly over the course of MG-132 treatment, consistent with impaired autophagic flux (Fig. 5C). The co-adaptor SINTBAD (also known as TBKBP1), which interacts with both TAX1BP1 and TBK1 during aggrephagy^9^, showed a similar pattern of increased stabilization in CDKL5 KO neurons, further supporting impaired autophagic clearance in the absence of CDKL5 (Fig. 5C). Consistent with this, treatment of neurons with the lysosomal inhibitor BafA1 increased levels of phosphorylated TBK1 in WT neurons, but not in CDKL5 KO neurons (Fig. 5D-E), indicating impaired autophagy-dependent turnover of activated TBK1 in the absence of CDKL5.

**Figure 5.**
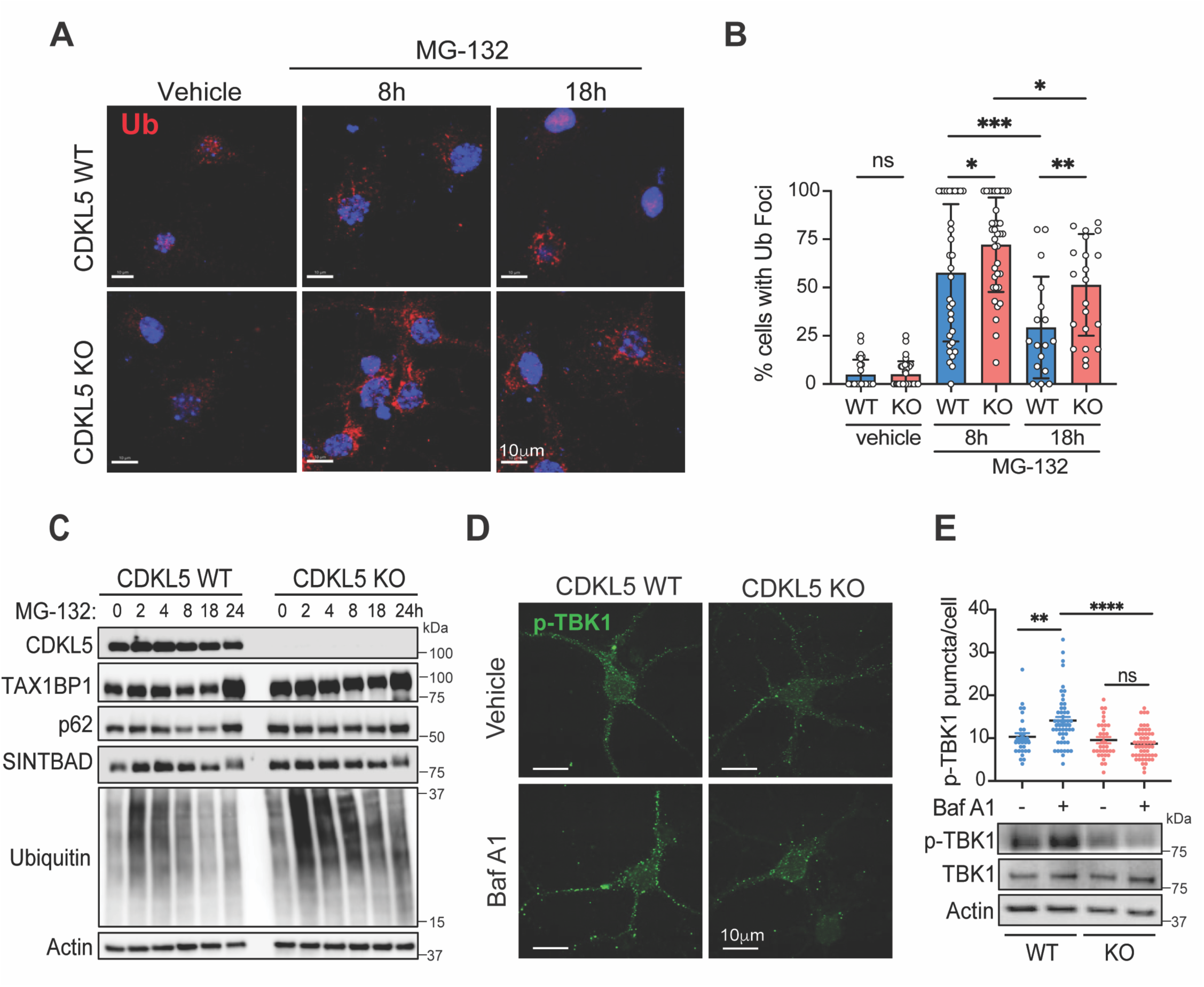
CDKL5-deficient primary cortical neurons have dysfunctional aggrephagy regulators. A) Representative fluorescence micrographs of mouse primary cortical neurons isolated from littermate CDKL5 WT and KO E16 embryos, and B) quantification of the percentage of neurons with ubiquitin foci. Bars represent the mean ± SE from 15-30 images, totaling at least 150 neurons counted across 3 independent embryos per genotype. C) Representative western blot of CDKL5 WT and KO cortical neurons. D) Representative fluorescence micrographs of primary cortical neurons treated with vehicle (DMSO) or BafA1 (500 nM) for 6 hours, with E) quantification of p-TBK1 puncta per cell (over 100 neurons quantified across 3 independent embryos per genotype) and a representative western blot of p-TBK1, total TBK1, and actin. Statistical analysis for B) and E) was performed using one-way ANOVA with Sidak’s correction for multiple comparisons. *p<0.05, **p < 0.005, ***p < 0.0005, ****p < 0.00005.

### CDKL5 regulates SINTBAD phosphorylation

Given that CDKL5 deficiency altered the turnover of SINTBAD and TAX1BP1, together with defective TBK1 activation, we hypothesized that SINTBAD, which binds both TBK1 and TAX1BP1, may be directly regulated by CDKL5. Using phosphorylation-site data deposited in the PhosphoSitePlus database, we searched SINTBAD for candidate phosphorylation sites conforming to the CDKL5 consensus motif (R-P-X-S/T-P/A). We identified Ser504, within the proline-rich region of SINTBAD, as a conserved candidate CDKL5 phosphorylation site (Fig. 6A). To determine whether CDKL5 interacts with SINTBAD, we performed co-immunoprecipitation in HeLa cells expressing HA-tagged SINTBAD. Immunoprecipitation of endogenous CDKL5 pulled down SINTBAD, and, in the reciprocal experiment, immunoprecipitation of SINTBAD pulled down CDKL5, indicating that the two proteins interact (Fig. 6B). Immunofluorescence colocalization studies showed that, at baseline, both SINTBAD and CDKL5 were diffusely distributed throughout the cytoplasm. Upon proteasome inhibition, both proteins redistributed into colocalized perinuclear puncta as early as 2 hours, before returning to a diffuse distribution by 16 hours (Fig. 6C).

**Figure 6.**
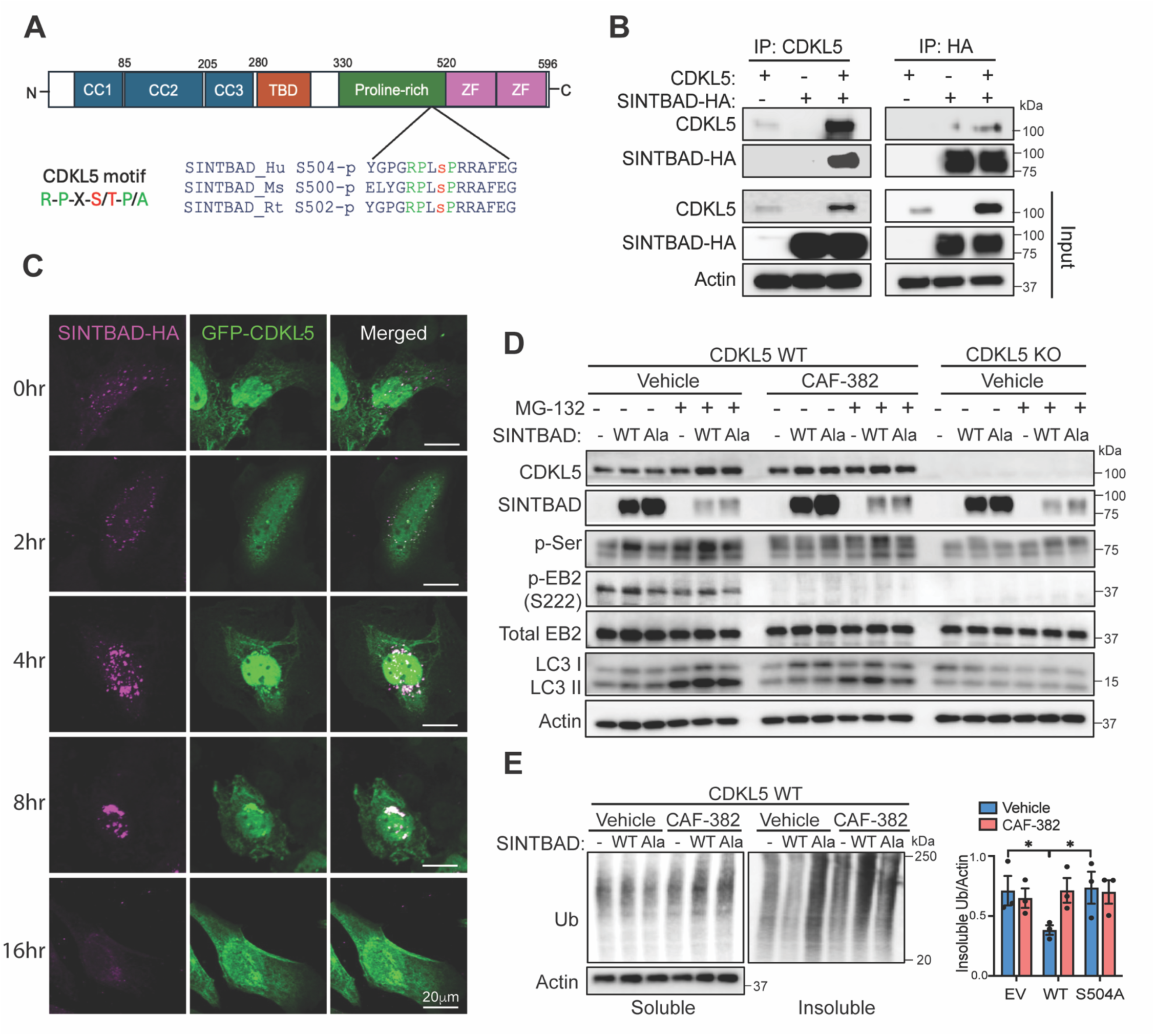
CDKL5 regulates SINTBAD phosphorylation and aggrephagy function. A) SINTBAD domain structure, highlighting the conserved Ser504 phosphorylation site, which conforms to the CDKL5 phosphorylation consensus motif. B) CDKL5 KO HeLa cells stably expressing EV or WT CDKL5 were transiently transfected with HA-tagged SINTBAD, followed by reciprocal immunoprecipitation with anti-CDKL5 or anti-HA antibodies and immunoblot detection of CDKL5, HA, and actin. C) Representative fluorescence micrographs of CDKL5 KO HeLa cells stably expressing GFP-CDKL5 and transfected with SINTBAD-HA, followed by treatment with MG-132 (1 μM) over a time course. D) Representative immunoblots of WT or CDKL5 KO HeLa cells transiently expressing SINTBAD-HA or the phospho-deficient S504A SINTBAD-HA mutant. WT HeLa cells were pretreated for 1 hour, and throughout the subsequent 8 hours of MG-132 treatment, with either DMSO or the CDKL5 inhibitor CAF-382. E) Lysates from WT HeLa cells overexpressing SINTBAD-HA or S504A SINTBAD-HA treated with DMSO or CAF-382 for 18 hours coupled with MG-132 were fractionated into Triton-soluble and - insoluble fractions and subjected to western blot analysis of ubiquitinated proteins.

To determine whether CDKL5 phosphorylates SINTBAD at Ser504, WT HeLa cells expressing empty vector (EV), WT SINTBAD, or a phospho-deficient SINTBAD mutant (S504A) were treated with the selective CDKL5 inhibitor CAF-382 plus MG-132, or vehicle plus MG-132 for 8 hours. SINTBAD phosphorylation was then assessed using a pan-phospho-serine antibody. CDKL5 KO cells were included as a control, and phosphorylation of EB2 (p-EB2) was used as a surrogate marker of CDKL5 activity to confirm loss of its kinase activity. Expression of WT SINTBAD increased serine phosphorylation signal at baseline, and this phosphorylation was further induced by MG-132 treatment; this MG-132-induced increase was not observed with the S504A mutant (Fig. 6D). This increase in WT SINTBAD phosphorylation was accompanied by increased LC3-II lipidation, suggesting that SINTBAD phosphorylation at Ser504 augments aggrephagy (Fig. 6D). Consistent with this, CDKL5 inhibitor treatment reduced both SINTBAD phosphorylation and LC3-II lipidation, and both were further reduced in CDKL5 KO cells, indicating that overexpression of WT SINTBAD is insufficient to rescue aggrephagy in the absence of CDKL5 activity (Fig. 6D).

To further support these findings, we assessed the accumulation of insoluble ubiquitinated proteins following MG-132 treatment, in the presence or absence of the CDKL5 inhibitor. Expression of WT SINTBAD suppressed the accumulation of insoluble ubiquitinated protein, whereas the S504A mutant failed to do so; however, this suppressive effect of WT SINTBAD was lost when CDKL5 activity was inhibited (Fig. 6E). Taken together, these findings suggest that CDKL5 phosphorylates SINTBAD at Ser504 to promote aggrephagy in response to proteasome inhibition.

### SINTBAD requires CDKL5 to promote the function of autophagy adaptors

To investigate the mechanism by which SINTBAD modulates aggrephagy, we expressed WT SINTBAD in HeLa cells in the presence or absence of the CDKL5 inhibitor CAF-382^20^. Following MG-132 treatment, TBK1 activation, assessed by Ser172 phosphorylation, increased most robustly in cells overexpressing SINTBAD, relative to SINTBAD-overexpressing cells treated with the CDKL5 inhibitor or to cells expressing an empty vector (Fig. 7A). SINTBAD overexpression also increased LC3-II levels and phosphorylation of p62 at Ser403 (Fig. 7A), a TBK1-dependent phosphorylation event that facilitates p62 binding to ubiquitinated proteins^10,24^. To validate these findings, WT and CDKL5 KO cells overexpressing SINTBAD were treated with puromycin, and protein levels were assessed over a 6-hour chase period following puromycin washout. SINTBAD overexpression promoted robust phosphorylation of TBK1 at Ser172 and p62 at Ser403, an effect that was most pronounced with BafA1 treatment. Additionally, following BafA1 treatment, levels of TAX1BP1, p62, and LC3-II accumulated to a greater extent in WT cells than in CDKL5 KO cells overexpressing SINTBAD, indicating that autophagic flux was more robustly induced in the presence of CDKL5 (Fig. 7B). Further investigation of the interaction between TAX1BP1 and SINTBAD by immunofluorescence revealed that the number of TAX1BP1/SINTBAD-positive foci was similar between WT and CDKL5 KO cells both at baseline and after the 6-hour chase period following puromycin treatment. However, BafA1 treatment significantly increased the number of these colocalized puncta in WT cells compared to CDKL5 KO cells (Fig. 7C-D). Together, these findings suggest that CDKL5 facilitates the co-adaptor function of SINTBAD, thereby enabling improved turnover of the aggrephagy adaptor proteins TAX1BP1 and p62, as well as activation of TBK1.

**Figure 7.**
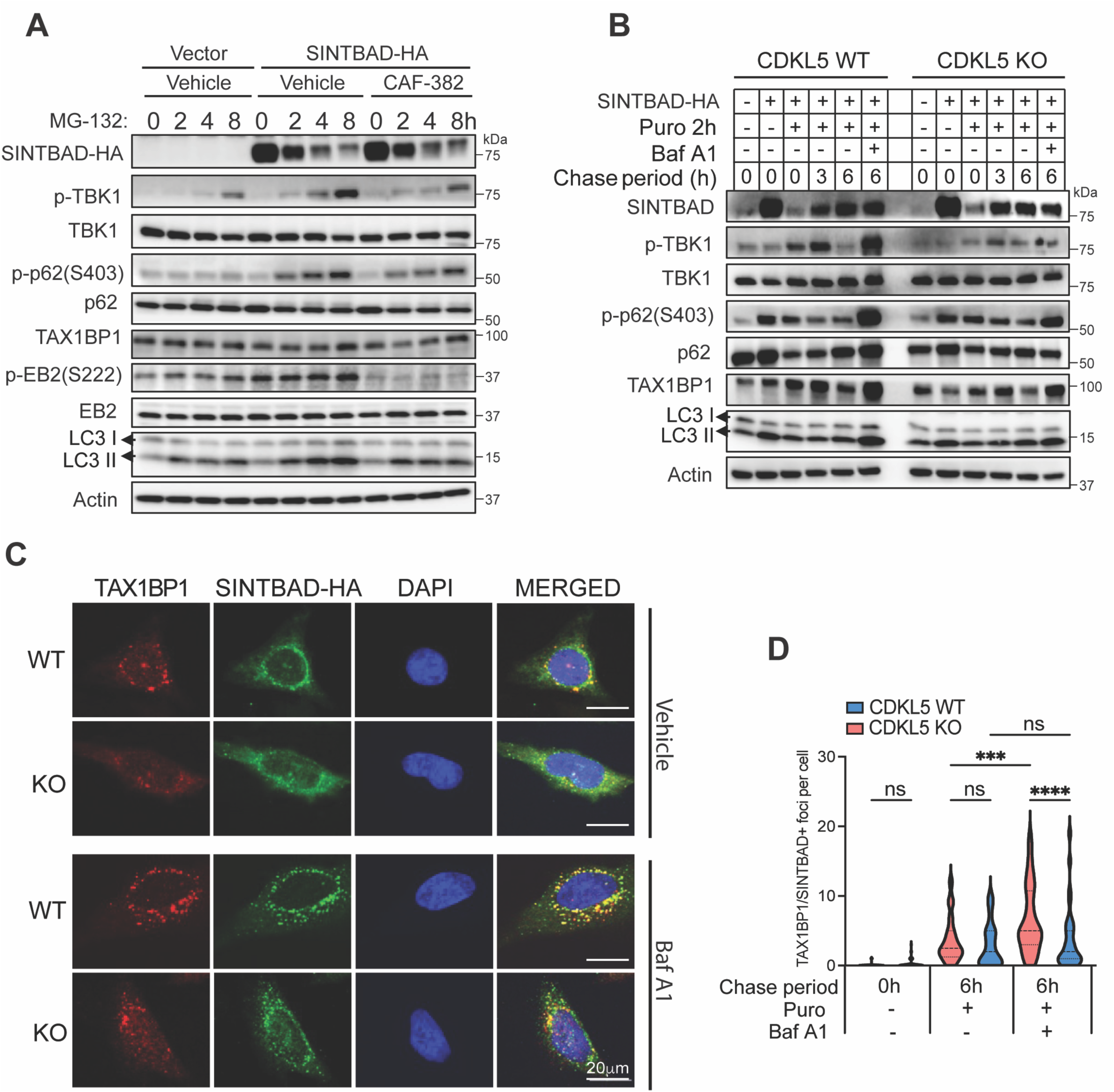
Overexpression of SINTBAD boosts aggrephagy in a CDKL5-dependent manner. A) Representative immunoblots of WT HeLa cells expressing empty vector or SINTBAD-HA, treated with DMSO or CAF-382, together with MG-132, for the indicated time points. Representative of three independent experiments. B-C) CDKL5 WT and KO HeLa cells expressing SINTBAD-HA, were treated with puromycin followed by a 6-hour chase period with or without BafA1 treatment for the final 4 hours. B) Representative immunoblot of three independent experiments. C) Representative fluorescence micrographs of endogenous TAX1BP1 and SINTBAD-HA in cells 6h post puromycin washout and treated with vehicle (DMSO) or BafA1. D) quantification of TAX1BP1-positive SINTBAD foci per cell. Histograms represent over 50 cells per condition across 3 independent experiments. Statistical analysis was performed using two-way ANOVA with Tukey’s correction for multiple comparisons. ***p < 0.0005, ****p < 0.00005.

We next investigated the necessity of SINTBAD phosphorylation at Ser504 during aggrephagy. Given that we observed early differences in TBK1 phosphorylation in the absence of CDKL5, we expressed WT, phospho-deficient (S504A), and phosphomimetic (S504E) SINTBAD constructs in CDKL5 KO cells and monitored colocalization between SINTBAD and phosphorylated TBK1 during early time points of MG-132 treatment (0-4hrs). The phospho-deficient SINTBAD mutant showed significantly fewer colocalized SINTBAD/TBK1 puncta at baseline compared to cells expressing WT or phosphomimetic SINTBAD. Notably, expression of phosphomimetic SINTBAD resulted in a significant increase in colocalized puncta after 2 hours of MG-132 treatment compared to WT SINTBAD, and after 2 and 4 hours compared to phospho-deficient SINTBAD (Fig. 8A-B). Consistent with the puromycin chase experiments in Figure 7, expression of SINTBAD S504E increased phospho-TBK1, phospho-p62, and TAX1BP1 levels after 6 hours of chase during autophagic flux inhibition with BafA1 (Fig. 8C).

**Figure 8.**
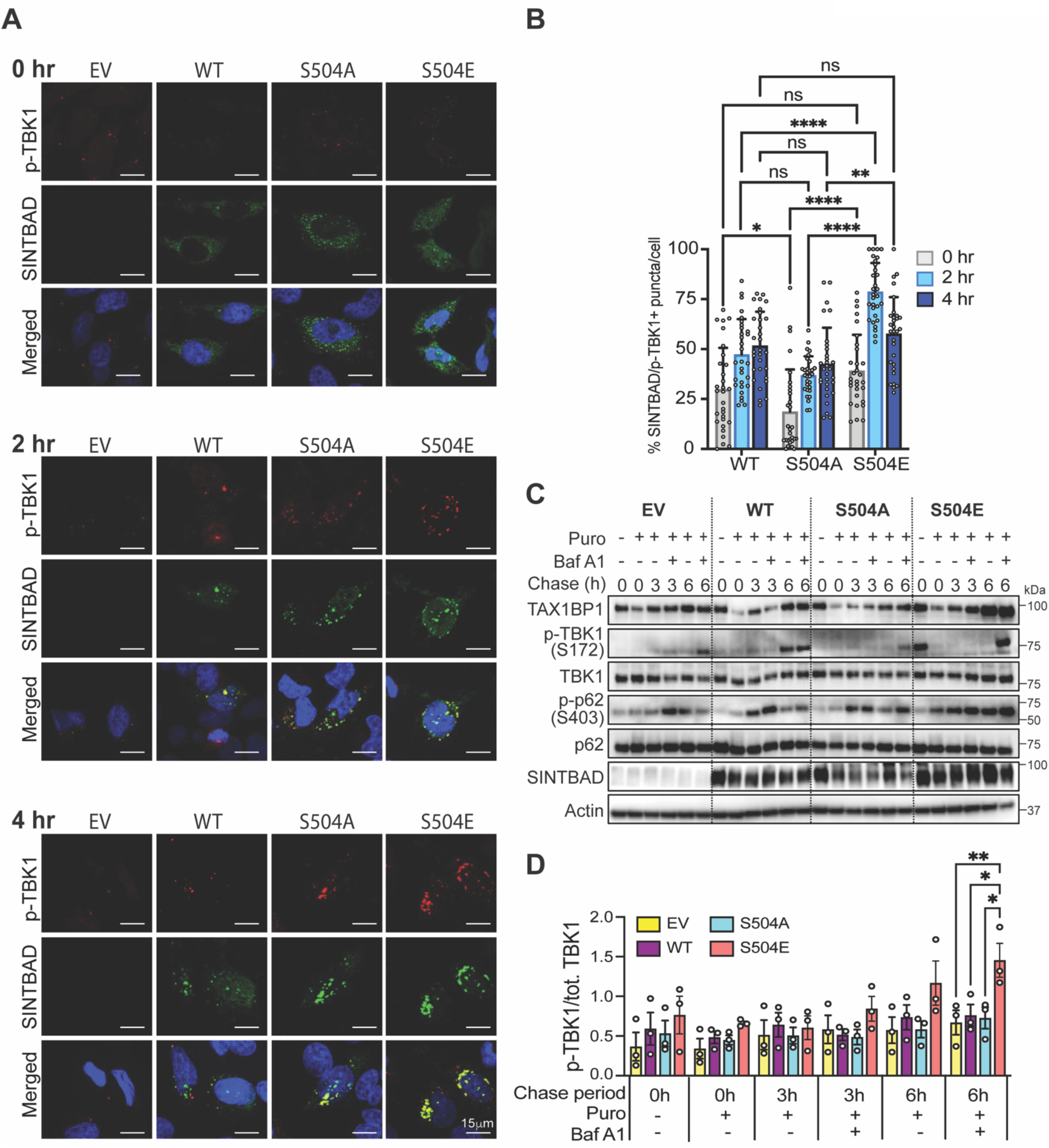
Phosphomimetic SINTBAD rescues pTBK1 phosphorylation. A) CDKL5 KO HeLa cells expressing empty vector, WT SINTBAD-HA, phospho-deficient SINTBAD-HA (S504A), or phosphomimetic SINTBAD-HA (S504E) were treated with MG-132 over a time course, with representative fluorescence micrographs of phospho-TBK1 (Ser172) and SINTBAD-HA, the latter detected using an anti-HA antibody. B) Quantification of the percentage of SINTBAD-positive TBK1 puncta per cell; 30 cells analyzed per condition; data representative of three independent experiments. Statistical analysis was performed using two-way ANOVA with Tukey’s correction for multiple comparisons. C) Immunoblots of CDKL5 KO HeLa cells expressing empty vector, WT SINTBAD-HA, phospho-deficient S504A SINTBAD-HA, or phosphomimetic S504E SINTBAD-HA, treated with puromycin followed by a 6-hour chase period in the presence or absence of BafA1. Blots representative of 3 independent experiments. D) Densitometry analysis of LC3-II normalized to actin. Bars represent the mean ± SE of three independent experiments. Statistical analysis was performed using two-way ANOVA with Sidak’s correction for multiple comparisons. *p<0.05, **p < 0.005, ****p < 0.00005.

Taken together, these findings indicate that CDKL5 mediates SINTBAD phosphorylation at Ser504, and that this phosphorylation event is required to drive SINTBAD’s co-adaptor function, promoting TBK1 activation, downstream phosphorylation of p62, and the coordinated turnover of TAX1BP1 and p62 during aggrephagy. In the absence of CDKL5, this regulatory axis is disrupted, resulting in impaired autophagic flux and defective clearance of ubiquitinated protein aggregates.

## Discussion

Maintenance of proteostasis requires cells to recognize misfolded, aggregated proteins and efficiently couple them to autophagy machinery for lysosomal degradation, a process called aggrephagy^7^. Here, we identify the neurodevelopmental kinase CDKL5 as a regulator of aggrephagy. Loss of CDKL5 impaired clearance of multiple classes of aggregation-prone proteins, including puromycin-induced aggregates, proteins accumulating during proteasome inhibition, and expanded polyglutamine-containing huntingtin, in both cell lines and primary neurons. This defect was also evident in vivo, where CDKL5-deficient mouse brains showed progressive, age-dependent accumulation of detergent-insoluble ubiquitinated proteins. Mechanistically, we place CDKL5 upstream of a SINTBAD–TBK1 signaling axis that coordinates the selective autophagy receptors TAX1BP1 and p62. We identify Ser504 of SINTBAD as a CDKL5-dependent phosphorylation site that promotes TBK1 activation, p62 Ser403 phosphorylation, TAX1BP1 recruitment, and clearance of ubiquitinated aggregates.

These findings extend our earlier work identifying CDKL5 as a regulator of p62-dependent virophagy, in which CDKL5 phosphorylates p62 at Thr269/Ser272 to promote capture of viral capsids^14^. The present results show that the role of CDKL5 in proteostasis is not restricted to viral cargo, but instead extends across mechanistically distinct forms of proteotoxic stress. The age-dependent aggregate phenotype we observe in CDKL5-deficient brains further supports this broader role in protein quality control, and raises the question of precisely where in the aggrephagy pathway CDKL5 acts.

To address this, we examined the recruitment of individual selective autophagy receptors to ubiquitinated cargo. In cell culture, p62 was recruited normally to ubiquitinated aggregates in CDKL5-deficient cells, but TAX1BP1 recruitment was markedly impaired, along with reduced TBK1 Ser172 and p62 Ser403 phosphorylation and diminished LC3-II accumulation. A similar pattern was evident in vivo: insoluble protein aggregates isolated from aged CDKL5 KO brains showed accumulation of p62, but proportionately less phospho-p62, alongside decreased recruitment of TAX1BP1. This pattern, in which p62 accumulates within aggregates without being efficiently phosphorylated or turned over, mirrors observations in human neurodegenerative proteinopathies and experimental models in which persistent aggregates remain p62-enriched but fail to recruit productive downstream autophagy machinery^29–32^.

Together, these results are consistent with an emerging model in which p62/NBR1 first condense ubiquitinated cargo, after which recruited TAX1BP1 couples these condensates to the autophagy initiation machinery and recruits TBK1 via SINTBAD and/or NAP1^8,9^. Within this model, our data suggest that CDKL5 facilitates the transition from initial cargo condensation to productive autophagic sequestration, rather than the initial cargo recognition step itself. Because TBK1-mediated p62 Ser403 phosphorylation enhances both ubiquitin binding and condensate degradation^10,11^, the impaired TBK1 activation in CDKL5 deficiency offers a mechanistic explanation for this impaired progression from condensate to degraded cargo.

SINTBAD, a TBK1 adaptor that links TAX1BP1 to TBK1 during aggrephagy, provides a likely connection in this pathway^9^. We found CDKL5 interacts with SINTBAD, and both transiently colocalize at perinuclear puncta during proteotoxic stress. SINTBAD phosphorylation increases in a CDKL5-and Ser504-dependent manner. Functionally, WT SINTBAD enhanced TBK1 activation, p62 phosphorylation, LC3 lipidation, and aggregate clearance, while the phospho-deficient S504A mutant did not. Conversely, phosphomimetic S504E enhanced TBK1 interaction and partially restored these responses in CDKL5-deficient cells. Together, these results support a model in which Ser504 phosphorylation enables SINTBAD to organize or sustain active TBK1 at sites of protein aggregation, potentially by concentrating TBK1 sufficiently to promote trans-autophosphorylation^25^.

Several questions remain regarding the broader scope of this regulatory mechanism. It is unclear whether CDKL5 similarly regulates the related TBK1 adaptor NAP1, which notably lacks a CDKL5 consensus motif^33^, or how this newly identified SINTBAD phosphorylation event relates to our previously described CDKL5-dependent phosphorylation of p62^14^. These open questions suggest that CDKL5 may coordinate multiple phosphorylation events that collectively facilitate autophagy activation, rather than acting through a simple one-kinase-one-substrate model. Further biochemical work, including in vitro kinase assays with purified CDKL5 and SINTBAD, will be needed to formally establish SINTBAD as a direct CDKL5 substrate and to determine how Ser504 phosphorylation affects its interactions with TBK1 and TAX1BP1.

This CDKL5–SINTBAD–TBK1 axis may be especially relevant in neurons, long-lived postmitotic cells that cannot dilute damaged and aggregation-prone proteins through cell division and are highly dependent on autophagic quality control^4^. TAX1BP1 is highly expressed in the brain and is required for efficient neuronal clearance of aggregation-prone proteins, including polyglutamine-expanded huntingtin^17^, consistent with the delayed turnover of ubiquitinated proteins we observed in CDKL5-deficient neurons. The progressive aggregate accumulation in aged CDKL5-deficient brains suggests that modest defects in this pathway may compound over a neuron’s lifetime, raising the possibility that impaired proteostasis contributes to the neurological dysfunction seen in CDKL5 deficiency disorder.

In summary, our findings identify CDKL5 as a regulator of aggrephagy that links proteotoxic stress to the TAX1BP1–SINTBAD–TBK1 selective-autophagy machinery, establishing a role for CDKL5 in long-term proteostasis and revealing a previously unrecognized regulator that facilitates the transition from protein aggregation to autophagic clearance.

## Materials and Methods

### Cell culture and reagents

HeLa cells were obtained from the American Type Culture Collection (ATCC) and cultured in Opti-MEM I Reduced Serum Medium supplemented with 5% FBS, 100 units/mL penicillin, and 100 µg/mL streptomycin (Thermo Fisher Scientific). Cell line identity was authenticated by ATCC. CDKL5 knockout (KO) HeLa clones, generated previously by CRISPR/Cas9-mediated disruption of exon 2 (Genome Engineering and iPSC Center, Washington University School of Medicine), were used throughout this study^14^. To generate cells stably expressing WT, KD or EV, CDKL5 KO HeLa cells were transduced with lentiviruses generated through the pLenti-C-Myc-DDK-IRES-Neo vector (Origene) as we previously described^14^. The selective CDKL5 kinase inhibitor CAF-382 was used at concentrations ranging from 10–100 nM, as indicated in figure legends, with an equivalent volume of DMSO used as vehicle control. MG-132 (EMD, 474791) was used at 0.1-1 µM for proteasome inhibition experiments.

### Primary cortical neuron culture

Cortical neurons were prepared from embryonic day 15-16 (E15-16) littermates generated by mating CDKL5+/-or CDKL5+/y mice. Cortices were dissected, dissociated in 2.5% trypsin (Thermo Fisher Scientific), and plated on glass chamber slides or 12-well plates pre-coated with 0.1% polyethyleneimine (Sigma-Aldrich). Cultures were maintained in Neurobasal medium supplemented with B27 (Invitrogen) and 2 mM glutamine; cytarabine (5 µM) was included for the first 24 hours to suppress proliferation of non-neuronal cells. Neurons were used for experiments after 7 days of differentiation.

### Mouse strains and aged brain tissue collection

CDKL5 KO and C57BL/6J wild-type (WT) mice were obtained from The Jackson Laboratory, with CDKL5 KO animals backcrossed for more than ten generations onto the C57BL/6J background. For analysis of age-dependent aggregate accumulation, WT and CDKL5 KO littermates were aged to the time points indicated in figure legends prior to euthanasia and brain collection. All animals were housed under pathogen-free conditions with a 12-hour light/dark cycle and ad libitum access to food and water. All procedures were approved by the University of Texas Southwestern Medical Center Institutional Animal Care and Use Committee.

### Assessment of protein aggregation in aged mouse brain

To assess age-dependent protein aggregate accumulation, brains were collected from WT and CDKL5 KO mice at 4, 18, and 24 months of age (n = 4–5 mice per genotype per age group). Brains were fixed in 4% paraformaldehyde, and frozen sections were cut at 5 µm thickness. Sections were stained with the PROTEOSTAT Aggresome Detection Kit (Enzo, ENZ-51035-K100) according to the manufacturer’s instructions to detect and quantify insoluble protein aggregates.

### Detergent soluble and insoluble protein fractionation

Brain tissue was homogenized (10% w/v) in Triton lysis buffer (50 mM Tris-HCl, pH 7.4, 150 mM NaCl, 1% Triton X-100, 0.5 mM DTT, supplemented with 1X protease inhibitor and 1X phosphatase inhibitor cocktails) and incubated on ice for 30 minutes. Homogenates were centrifuged at 3,000 × g for 10 minutes at 4°C to pellet debris. The resulting supernatant was transferred to a new tube and centrifuged at 20,000 × g for 30 minutes at 4°C; this second supernatant was retained as the detergent-soluble fraction. The pellet was washed twice with 1 mL of lysis buffer, with re-pelleting by centrifugation at 20,000 × g between washes and then resolubilized in lysis buffer to yield the detergent-insoluble fraction. Both fractions were combined with 2X Laemmli sample buffer and boiled for 5 minutes prior to SDS-PAGE and western blot analysis of ubiquitinated protein, p62, TAX1BP1, and SINTBAD content. The same fractionation approach was applied to cultured cells, substituting cell pellets for brain tissue.

### Generation of SINTBAD constructs

The HA-tagged SINTBAD backbone vector, pBMN-HA-SINTBAD (Addgene, plasmid #210209), was used to generate phospho-deficient (S504A) and phosphomimetic (S504E) point mutants. Site-directed mutagenesis was performed using the QuikChange II Site-Directed Mutagenesis Kit (Agilent) according to the manufacturer’s instructions. For the S504A mutant, the following primer pair was used: forward, 5’-GGCGCGCCGCGGTGCGAGGGGCCTGCC-3’; reverse, 5’-GGCAGGCCCCTCGCACCGCGGCGCGCC-3’. For the S504E mutant, the primers used were: forward, 5’-GGCGCGCCGCGGTTCGAGGGGCCTGCC-3’; reverse, 5’-GGCAGGCCCCTCGAACCGCGGCGCGCC-3’. All mutant constructs were verified by Sanger sequencing prior to use in downstream experiments. WT and mutant SINTBAD constructs, along with empty vector control, were introduced into HeLa cells (parental or CDKL5 KO) by transient transfection.

### Induction and chase of protein aggregates

To induce aggresome-like inclusions, cells were treated with puromycin (Gibco, A11138-03; 5 µg/mL) for 2 hours, after which puromycin-containing media was removed and replaced with fresh culture media; cells were then harvested or fixed at the time points indicated in figure legends to monitor clearance of ubiquitin-positive foci during the chase period. For proteasome inhibition experiments, cells were treated with MG-132 (EMD, 474791; 0.1-1µM) for the durations indicated in figure legends. Autophagic flux was assessed in parallel by co-treatment with Bafilomycin A1 (BafA1, 200 nM). For polyglutamine aggregation studies, HeLa cells were transfected with constructs encoding an expanded polyglutamine repeat, pEGFPQ-23 (Addgene; plasmid #40261) and pEGFPQ-74 (Addgene; plasmid #40262), for 48 hours; cell lysates were fractionated by centrifugation at 500 × g into soluble and insoluble fractions and subjected to filter trap assay as previously described^34^. Detection of huntingtin was performed using an anti-GFP antibody.

### Co-immunoprecipitation

To assess the interaction between CDKL5 and SINTBAD, HeLa cells expressing HA-tagged SINTBAD were washed with cold 1X PBS, scraped, and pelleted by centrifugation at 400 × g for 2 minutes. Cell pellets were lysed in 1X RIPA buffer (Cell Signaling Technology) for 15 minutes on ice, and lysates were clarified by centrifugation at 15,000 × g for 10 minutes at 4°C. Clarified lysates were incubated overnight at 4°C with anti-HA magnetic beads (MBL, HA-tagged Protein Purification Kit, catalog #3320). Beads were washed five times with lysis buffer, and bound proteins were eluted by boiling in 2X Laemmli sample buffer for 5 minutes prior to SDS-PAGE and western blot analysis.

### Antibodies

The following primary antibodies were used for western blot (WB) and/or immunofluorescence (IF): rabbit anti-CDKL5 (EMD, MABS1132; WB), mouse anti-CDKL5 (Coriell, OR00047 and OR00049; WB), mouse anti-ubiquitin (Santa Cruz, SC-9133, WB/IF; SC-8017, WB), rabbit anti-ubiquitin (Cell Signaling, 58395; WB/IF), rabbit anti-TAX1BP1 (Proteintech, 14424-1-AP; WB/IF), rabbit anti-TAX1BP1 (Invitrogen, 702840; WB), rabbit anti-phospho-p62 (Ser403) (Cell Signaling, 39786S; WB), rabbit anti-phospho-p62 (Thr269/Ser272) (Cell Signaling, 13121S; WB), mouse anti-p62/SQSTM1 (Abnova, H00008878-M01; WB/IF), guinea pig anti-p62/SQSTM1 (Progen, GP62-C; WB/IF), mouse anti-actin-HRP (Santa Cruz, SC-47778-HRP; WB), rabbit anti-TBK1 (Cell Signaling, 3504S; WB), rabbit anti-phospho-TBK1 (Ser172) (Cell Signaling, 5483S; WB/IF), mouse anti-puromycin (EMD, MABE343; WB), rabbit anti-LC3B (Novus Biologicals, NB100-2220; WB), rabbit anti-SINTBAD (Cell Signaling, 8605S; WB), rabbit anti-HA (Cell Signaling, 3724S; WB/IF), rabbit anti-EB2 (Abcam, ab45676; WB), chicken anti-GFP (Abcam, ab13970; WB) and rabbit anti-phospho-EB2 (Covalab, pAb01032-P; WB).

HRP-conjugated secondary antibodies used for western blot were: goat anti-mouse IgG (EMD, 115-035-071), donkey anti-rabbit IgG (EMD, AP182P), donkey anti-sheep IgG (EMD, 61-8620), goat anti-guinea pig IgG (Abcam, ab6908), and goat anti-chicken IgG (Abcam, ab6877). For immunofluorescence, the following Alexa Fluor-conjugated secondary antibodies were used (all Invitrogen): goat anti-mouse 488 (A21042), goat anti-mouse 568 (A11031), goat anti-mouse 594 (A11013), goat anti-mouse 647 (A21235), goat anti-rabbit 488 (A11008), goat anti-rabbit 594 (A11012), and goat anti-rabbit 647 (A21245).

### Immunofluorescence and colocalization analysis

HeLa cells and cortical neurons were cultured on glass chamber slides, fixed with 4% paraformaldehyde in PBS, and permeabilized with ice-cold methanol for 20 minutes. For CDKL5-GFP-expressing cells, CDKL5 KO cells were transfected with pEGFP-CDKL5 (Addgene, plasmid #245899). After blocking in PBS containing 3% BSA, cells were incubated with primary antibodies against ubiquitin, GFP-CDKL5 (anti-GFP), SINTBAD-HA (anti-HA), p62, TAX1BP1, or phospho-TBK1 (Ser172), followed by the Alexa Fluor-conjugated secondary antibodies listed above. Images were acquired on a Zeiss AxioImager Z2 microscope or a Nikon CSU-W1 spinning-disk confocal microscope. Colocalized puncta were quantified per cell using Imaris (v10.2.0). Nuclei were segmented by absolute intensity thresholding, and puncta were detected using the Spots algorithm (estimated diameter, 0.5 µm) with region growing and object-object statistics enabled. Spots within a distance of 0.5 µm or less were considered colocalized.

### Live-cell time-lapse imaging and analysis

WT and CDKL5 KO HeLa cells were transfected with a pBMN-mEGFP-TAX1BP1 plasmid (obtained from Richard Youle)^35^ for 24 hours prior to imaging. Time-lapse imaging was performed on a Nikon CSU-W1 spinning-disk confocal microscope equipped with Perfect Focus System and a Tokai environmental control incubator, using a 60x oil-immersion objective. Immediately before imaging, media was replaced with puromycin (1:2000), and 3D image stacks (0.75 µm step size; 6.0 µm total range) were acquired at four fields of view every 30 seconds for 2 hours.

Time-lapse data were analyzed in Imaris (v9.6.1; Oxford Instruments) following deconvolution using the software’s standard iterative algorithm (10 iterations), with parameters set for the acquisition system used. TAX1BP1-GFP puncta were detected in 10 WT and 10 KO cells using the Spots algorithm (estimated diameter, 0.6 µm) with region growing and tracking enabled; punctum fusion events were identified using the Connected Components algorithm, and spot counts were exported at each time point.

### Western blot analysis

Cells or tissue were lysed in RIPA buffer (Cell Signaling Technology, 9806) or Triton buffer (50 mM Tris, 1 mM EDTA, 150 mM NaCl, 1% Triton X-100) supplemented with protease and phosphatase inhibitor cocktails, and clarified by centrifugation at 10,000 × g for 10 minutes at 4°C, and the supernatants diluted with 2× Laemmli sample buffer (Bio-Rad Laboratories) containing 5% β-mercaptoethanol. Lysates were resolved on 4–20% gradient polyacrylamide gels (Bio-Rad) and transferred to PVDF membranes. Membranes were blocked in 5% non-fat milk in 1× TBS buffer containing 0.05% Tween 20 and probed with the primary and secondary antibodies listed above. Signal was detected using enhanced chemiluminescence (Thermo Fisher Scientific) and imaged on a Bio-Rad ChemiDoc system; band intensity was quantified using ImageJ2.

### Statistical analysis

All statistical analyses were performed using GraphPad Prism unless otherwise noted below. Two-group comparisons of normally distributed data were assessed by unpaired two-tailed t-tests.

Comparisons across more than two groups were performed by one-way ANOVA with Dunnett’s post hoc test, or two-way ANOVA where two independent variables were assessed jointly. All experiments were performed in at least three independent biological replicates, as indicated in figure legends. A p-value of < 0.05 was considered statistically significant.

Time-lapse spot-count data were analyzed separately in RStudio (v2024.04.2) using the R packages dplyr, tidyr, mcp, nlme, and emmeans. A Bayesian changepoint model was fit to WT-KO difference scores to identify the time point at which clearance kinetics diverged between genotypes. Prior to this changepoint, natural log-transformed spot counts were analyzed using a linear mixed-effects model with a compound symmetry, heteroscedastic variance structure; estimated marginal means were compared at 5-minute intervals, with t-statistics and p-values derived using Satterthwaite-approximated degrees of freedom and adjusted for multiple comparisons using the Hommel method.

## Acknowledgements

This work was supported by NIH KO8AI163377, Burroughs Wellcome Fund 1022205, and the Disease Oriented Clinical Scholars Program at UT Southwestern. The authors would like to acknowledge the Quantitative Light Microscopy Core, a Shared Resource of the Harold C. Simmons Comprehensive Cancer Center, supported in part by an NCI Cancer Center Support Grant, 1P30 CA142543-01. We would additionally like to acknowledge the UT Southwestern Whole Brain Microscopy and Neuro-Models Facilities for their assistance with mouse brain imaging. We thank Alison Axtman for providing us with the CDKL5 inhibitor, CAF-382, and Richard Youle for the TAX1BP1-expressing vector.

## Author contributions

JT conceptualized and supervised the study. JT, ZZ and BK analyzed the data. JT acquired funding. ZZ, BK, SS, JL, and GR carried out the investigation. JT wrote the original draft of the manuscript, which was reviewed and edited by all authors.

