## Supplemental Figures for "CDKL5 deficiency impairs TBK1-mediated autophagy and clearance of neuronal protein aggregates"

### Supplemental Figure 1

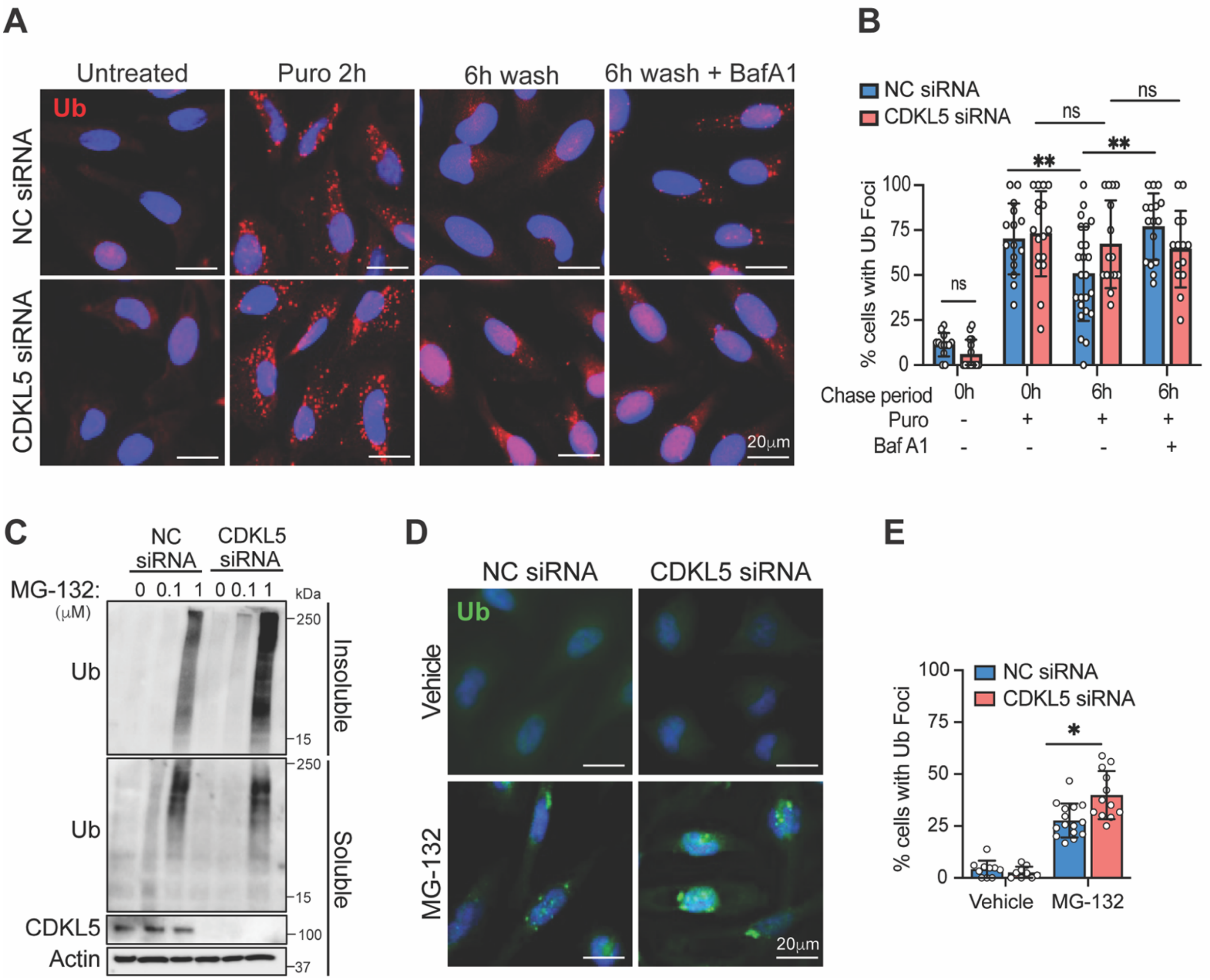

### Supplemental Figure 2

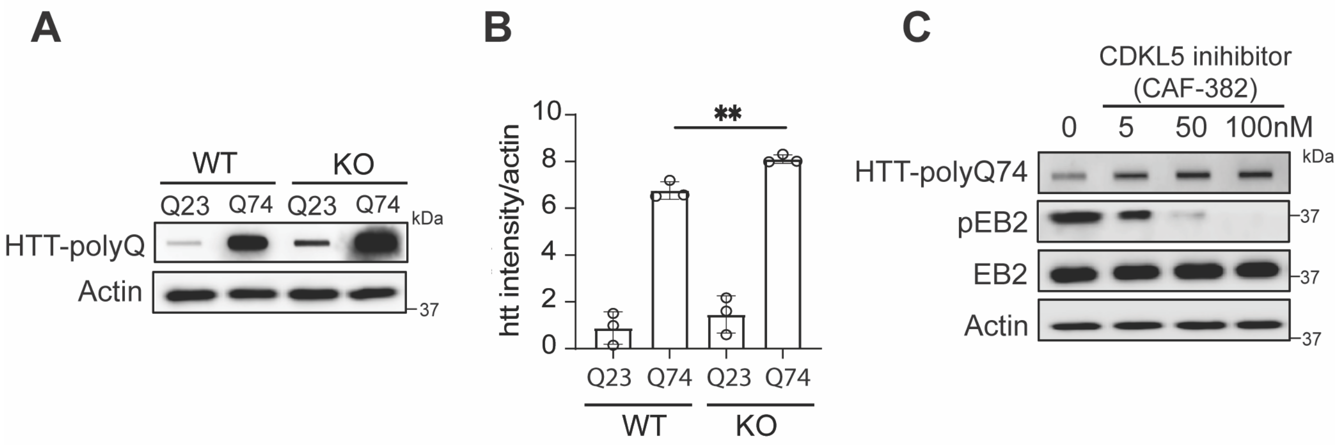

### Supplemental Figure 3

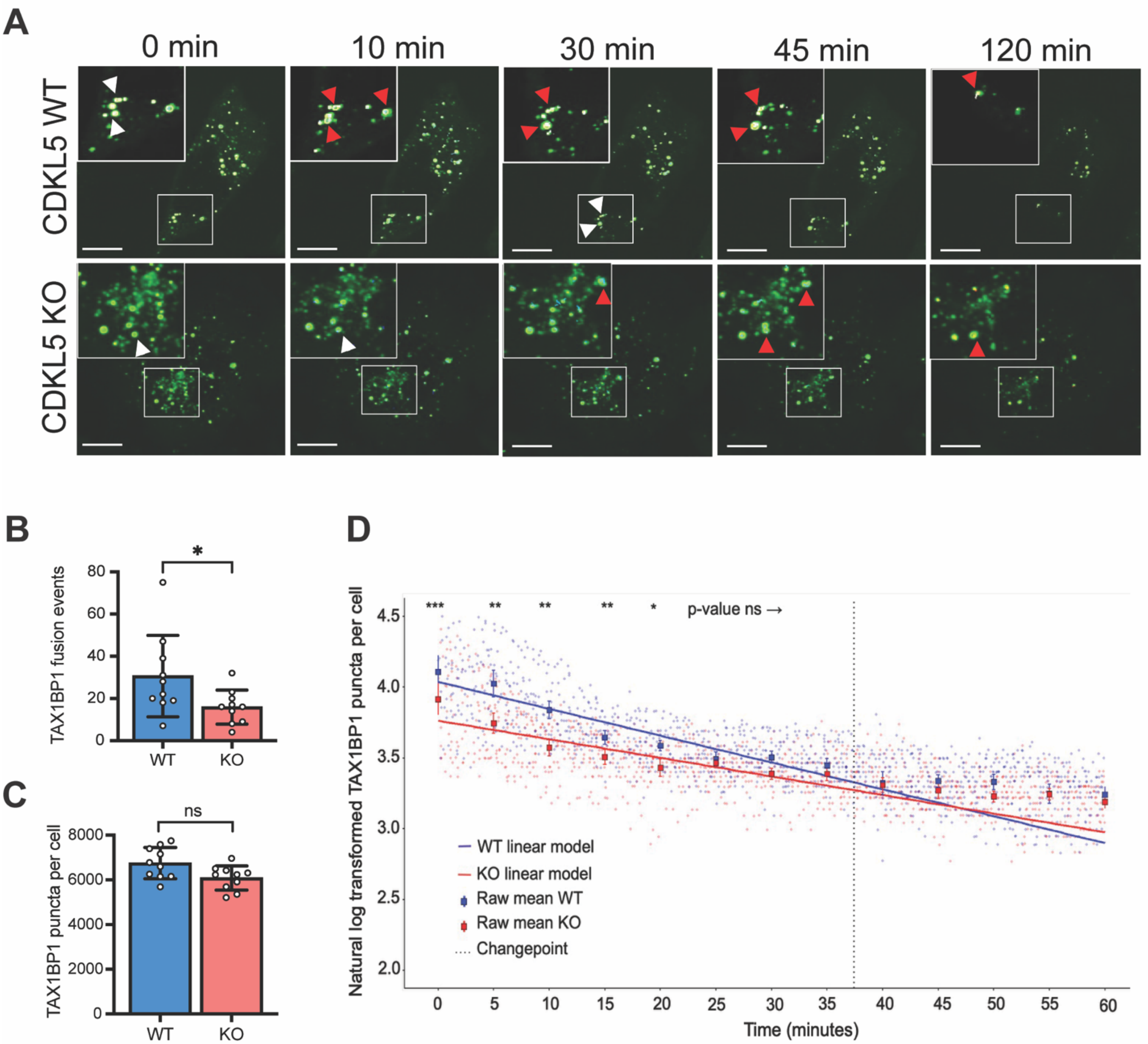

**Supplemental Figure Legends**

**Supplemental Figure 1. CDKL5 knockdown impairs clearance of protein aggregates.** A) Representative fluorescence micrographs of ubiquitin in HeLa cells subjected to non-coding or CDKL5-targeting siRNA, then treated with 5 μg/mL puromycin for 2 hours, washed, and replaced with fresh media containing BafA1 (200 nM) for the designated chase period, with B) quantification of the percentage of cells with ubiquitin foci from 15 images and over 150 cells per condition across 3 independent experiments. Histobars represent the mean ± SE. Statistical analysis was performed using one-way ANOVA with Sidak's correction for multiple comparisons. C) Lysates from HeLa cells treated with non-coding or CDKL5 siRNA and increasing concentrations of MG-132 for 18 hours were fractionated into Triton-soluble and -insoluble fractions for western blot analysis of ubiquitinated proteins, CDKL5 expression, and actin. D) Representative fluorescence micrographs and E) quantification of the percentage of cells with ubiquitin foci. Histobars represent the mean ± SE of 12 images and over 100 cells per condition across 3 independent experiments. Statistical analysis was performed using one-way ANOVA with Sidak's correction for multiple comparisons.

**Supplemental Figure 2. CDKL5 regulates clearance of polyglutamine-expanded huntingtin aggregates.** A) WT and CDKL5 KO HeLa cells expressing GFP-tagged HttQ23 or HttQ74 were subjected to filter trap assay, with detection of GFP-Htt in the trapped aggregate fraction and actin in the soluble fraction by western blot. B) Densitometry quantification of Htt aggregates normalized to actin from three independent filter trap experiments. Bars represent the mean ± SE. Statistical analysis was performed using one-way ANOVA with Sidak's correction for multiple comparisons. **$p<0.005$. C) HeLa cells expressing HttQ74 were treated with increasing concentrations of the CDKL5 inhibitor CAF-382, followed by filter trap assay with western blot detection of HttQ74 in the aggregate fraction. CDKL5 activity was assessed by phosphorylation of EB2 at Ser222.

**Supplemental Figure 3. TAX1BP1 fusion kinetics during proteotoxic stress are impaired in CDKL5 deficiency.** A) Representative time-lapse fluorescence micrographs of WT and CDKL5 KO cells expressing GFP-TAX1BP1, treated with puromycin and imaged every 30 seconds for 2 hours. White arrows indicate non-fused puncta that subsequently fuse, as marked by red arrows in later frames. B) Quantification of TAX1BP1 puncta fusion events per cell, detected using the Connected Components tracking algorithm. C) Cumulative number of detected TAX1BP1 puncta per cell across the time-lapse period. D) Linear mixed-effects model analysis of the kinetic change in TAX1BP1 puncta counts over time in WT and CDKL5 KO cells, with the estimated changepoint denoting the time point up to which WT and CDKL5 KO clearance kinetics are statistically divergent. Data are representative of 10 WT and 10 CDKL5 KO cells analyzed. Statistical analysis was performed using a linear mixed-effects model with a compound symmetry, heteroscedastic variance structure; estimated marginal means were compared using t-statistics with Satterthwaite-approximated degrees of freedom, and p-values were adjusted for multiple comparisons using the Hommel method.
